# Pluripotent Stem Cell-derived Cerebral Organoids Reveal Human Oligodendrogenesis with Dorsal and Ventral Origins

**DOI:** 10.1101/460907

**Authors:** Hyosung Kim, Ranjie Xu, Padmashri Ragunathan, Anna Dunaevsky, Ying Liu, Cheryl F. Dreyfus, Peng Jiang

**Affiliations:** Department of Cell Biology and Neuroscience, Rutgers University, Piscataway, NJ 08854, USA; Department of Developmental Neuroscience, Munroe-Meyer Institute, University of Nebraska Medical Center, Omaha, NE 68198, USA; Department of Neurosurgery and Center for Stem Cell and Regenerative Medicine, the Brown Foundation Institute of Molecular Medicine for the Prevention of Human Diseases, University of Texas Health Science Center at Houston, Houston, TX 77030, USA; Department of Neuroscience and Cell Biology, Rutgers Robert Wood Johnson Medical School, Piscataway, NJ 08854, USA

## Abstract

The process of oligodendrogenesis has been relatively well delineated in the rodent brain. However, it remains unknown whether analogous developmental processes are manifested in the human brain. Here, we report oligodendrogenesis in forebrain organoids, generated by using OLIG2-GFP knockin human pluripotent stem cell (hPSC) reporter lines. OLIG2/GFP exhibits distinct temporal expression patterns in ventral forebrain organoids (VFOs) vs. dorsal forebrain organoids (DFOs). Interestingly, oligodendrogenesis can be induced in both VFOs and DFOs after neuronal maturation. Assembling VFOs and DFOs to generate fused forebrain organoids (FFOs) promotes oligodendroglia maturation. Furthermore, dorsally-derived oligodendroglial cells outcompete ventrally-derived oligodendroglia and become dominant in FFOs after long-term culture. Thus, our organoid models reveal human oligodendrogenesis with ventral and dorsal origins. These models will serve to study the phenotypic and functional differences between human ventrally- and dorsally-derived oligodendroglia and to reveal mechanisms of diseases associated with cortical myelin defects.

## INTRODUCTION

Oligodendrocytes (OLs) are the myelinating cells and are the last major type of neural cells formed during the CNS development (Goldman and Kuypers, 2015; Zhong et al., 2018). Much of our current knowledge on OL development is obtained from studies in rodents. There are three distinct and sequential waves of oligodendrogenesis. Starting at embryonic day (E) 12.5, the first wave arises from NKX2.1-expressing precursors in the medial ganglionic eminence (Tekki-Kessaris et al., 2001). These oligodendrocyte progenitor cells (OPCs) migrate tangentially to distant locations and colonize the entire forebrain (Kessaris et al., 2006; Klambt, 2009). Subsequently, at E15.5, a second wave emerges from precursors in the lateral and medial ganglionic eminences (Chapman et al., 2013). These OPCs also migrate to the cortex, dispersing throughout the forebrain (Kessaris et al., 2006; Klambt, 2009). These ventrally-derived OPCs either remain as OPCs or differentiate into myelinating OLs in the maturing neocortex (Kessaris et al., 2006; Tripathi et al., 2011). Around birth, a third wave arises dorsally from precursors expressing the homeobox gene EMX1 in the cortex (Kessaris et al., 2006; Winkler et al., 2018). Interestingly, these dorsally-derived OPCs migrate locally to populate the cortex and displace the OPCs from the first two waves that initially colonize the cortical mantle (Kessaris et al., 2006; Winkler et al., 2018). Although the process of oligodendrogenesis has been well delineated in the mouse brain, it remains unknown whether analogous developmental processes are manifested in the human brain, particularly the dorsally-derived third wave of oligodendrogenesis (Jakovcevski et al., 2009; Rakic and Zecevic, 2003).

The lack of human brain tissue prevents a detailed understanding of human oligodendrogenesis. Neural specification of human pluripotent stem cell (hPSC), including human embryonic stem cells (hESCs) and human induced pluripotent stem cells (hiPSCs), offers unprecedented opportunities for studying human neural development (Avior et al., 2016; Marchetto et al., 2011). While OPCs have been efficiently derived from hPSC in the 2-dimensional (2D) culture system, these OPCs are mainly derived from ventral CNS regions by using ventralizing morphogens, such as sonic hedgehog (SHH) (Goldman and Kuypers, 2015; Tao and Zhang, 2016). Despite the previous progress, the different origins of human oligodendrogenesis have not been able to be recapitulated in 2D cultures, likely due to the lack of cell-cell/cell-matrix interactions in these monolayer cultures.

The recent development of 3D cerebral organoids derived from hPSCs offers a promising approach to understanding human brain development (Pasca, 2018), complementing 2D cell culture models. In this study, using OLIG2-GFP knockin hPSC reporter lines (Liu et al., 2011), we generated brain region-specific dorsal forebrain organoids (DFOs) and ventral forebrain organoids (VFOs) by inhibiting or activating SHH signaling pathway, respectively. We further monitored the temporal expression of OLIG2 and examined the oligodendrogenesis in these brain region-specific organoids. Moreover, by assembling DFOs and VFOs to form fused forebrain organoids (FFOs), we found that the human oligodendroglial differentiation and maturation were significantly promoted.

## RESULTS

### Distinct expression patterns of OLIG2 in hPSC-derived ventral and dorsal forebrain organoids

To examine the expression of OLIG2 in brain region-specific organoids, we generated human ventral forebrain organoids (VFOs) and dorsal forebrain organoids (DFOs) using OLIG2-GFP hPSC (hESC and hiPSC) reporter lines generated in our previous studies (Fig. 1A and B) (Liu et al., 2011; Xue et al., 2009). Using methods established in recent studies (Bagley et al., 2017; Birey et al., 2017; Xiang et al., 2017), brain region-specific organoids were derived by either activating or inhibiting the SHH signaling pathway from week 3 to 5. After one additional week of culturing with neural differentiation (ND) medium or OPC medium (week 6), both DFOs and VFOs exhibited subventricular/ventricular zone-like regions containing SOX2^+^ neural progenitor cells (NPCs) and βIIIT^+^ immature neurons (Fig. 1C). As shown in Fig. 1D and E, at week 5, both VFOs and DFOs were composed of a comparable population of Nestin-positive NPCs and FOXG1-positive forebrain cells. The vast majority of the cells in DFOs remained to express PAX6, which is also a marker for the NPCs in dorsal brain, whereas the vast majority of the cells in VFOs expressed NKX2.1, a marker of ventral prosencephalic progenitors. We further examined the gene expression of markers for dorsal forebrain, including *EMX1* and *TBR2*, and markers for ventral forebrain, including *NKX2.2*, *DLX1*, and *LHX6* in both DFOs and VFOs. EMX1 and TBR2 are expressed by cortical NPCs and intermediate progenitors (Englund et al., 2005; Gorski et al., 2002). NKX2.2, LHX6, and DLX1 are expressed by the NPCs in the medial ganglionic eminence (Briscoe et al., 1999; Du et al., 2008; Petryniak et al., 2007). As shown in Fig. 1F, markers for ventral forebrain, *NKX2.2*, *DLX1*, and *LHX6*, were conspicuously detected in VFOs, whereas markers for dorsal forebrain, *EMX1* and *TBR2*, were restricted to DFOs. Although DFOs and VFOs were generated from the same population of PAX6^+^ pNPCs at week 3 (Fig. 1B), as opposed to week 5 DFOs, nearly all the cells in week 5 VFOs were NKX2.1^+^/PAX6^−^ and abundantly expressed *NKX2.2*, *DLX1*, and *LHX6*, indicating that VFOs undergo ventral forebrain development induced by the activation of the SHH signaling pathway. Together with the data showing the restricted expression of *EMX1* and *TBR2* in DFOs, the observation that week 5 DFOs were highly enriched of PAX6^+^/NKX2.1^−^ NPCs indicates the formation of dorsal forebrain regional identity in DFOs. From week 5 to 7, intense GFP signals were observed in VFOs, whereas a small subset of cells in DFOs was found to express GFP (Fig. 1B and G). After long-term culture, robust GFP fluorescence in the VFOs became dimmer at week 9 and eventually was found to distribute evenly in the VFOs at week 12. The weak GFP signals in the DFOs gradually decreased and became undetectable at week 9. Interestingly, we observed the reappearance of GFP signals at week 12 (Fig. 1G). Furthermore, we confirmed the *OLIG2* expression in DFOs by qRT-PCR. We consistently found that *OLIG2* expression was very low at week 5 and hardly detectable at week 9. At week 12, the *OLIG2* expression significantly increased about 25 fold, compared to its level at week 5 (Fig. 1H).

**Figure 1.**
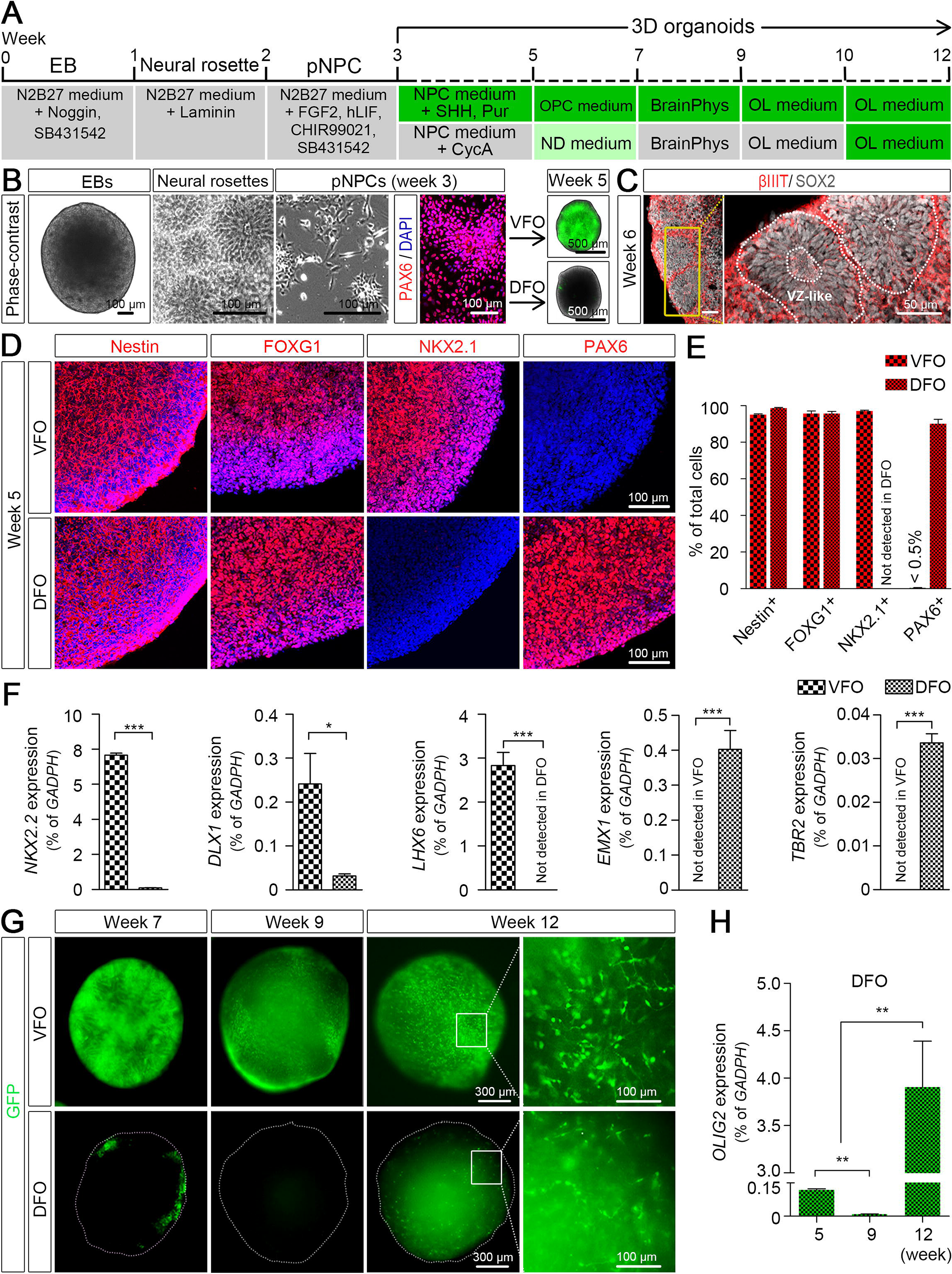
Temporal expression of OLIG2 in hPSC-derived VFOs and DFOs. (A) A schematic procedure for deriving brain region-specific forebrain organoids from OLIG2-GFP hPSCs by the treatment of a combination of sonic hedgehog (SHH) and purmorphamine (Pur) or cyclopamine (CycA) alone for VFOs and DFOs, respectively. The stages after week 3 are color-coded based on the expression of GFP. (B) Representative bright-field and fluorescence images of embryoid bodies (EBs) at week 1, neural rosettes at week 2, primitive neural progenitor cells (pNPCs) at week 3, and VFOs and DFOs at week 5. pNPCs at week 3 were positive for PAX6 staining. Scale bars: 100 μm for bright-field images and 500 μm for fluorescence images. (C) Representatives of the ventricular zone (VZ)-like structure formed by βIIIT and SOX2^+^ cells in FOs at week 6. Scale bars, 50 μm. (D and E) Representatives and quantification of Nestin-, FOXG1-, NKX2.1-, and PAX6-expressing cells in week 5 VFOs or DFOs (n = 4 organoids from two hPSC lines). (F) qRT-PCR results showing the expression of *NKX2.2*, *DLX1*, *LHX6*, *EMX1*, and *TBR2* in week 5 VFOs and DFOs (n = 3, independent experiments). Student’s *t*-test., **p < 0.05 and ***p < 0.001. (G) Temporal expression of GFP fluorescence in VFOs and DFOs. Scale bar: 300 μm in the original images and 100 μm in the enlarged images. (H) qRT-PCR results showing the expression of *OLIG2* at different time points in the DFOs. The expression level is normalized to GAPDH (n = 4, independent experiments). One-way ANOVA with Turkey’s post hoc test. **p < 0.01.

### OLIG2 is cytoplasmically expressed in PAX6^+^ neural progenitors in week 5 DFOs

GFP signals faithfully mirrored the OLIG2 expression in organoids (Fig. 2A and B). There was a significantly higher abundance of OLIG2^+^ cells in VFOs than in DFOs. Notably, unlike the nuclear localization of OLIG2 in VFOs, GFP^+^ cells in DFOs exhibited cytoplasmic OLIG2 expression (Fig. 2A and C). Immunoblot analysis confirmed that OLIG2 was abundantly present in the nuclear fraction of VFOs, whereas OLIG2 was detected at a low level only in the cytoplasmic fraction of DFOs (Fig. 2D). In the VFOs, nearly all GFP^+^ cells expressed NKX2.1 (Fig. 2E). As predicted, virtually no detectable NKX2.1^+^ cells in the DFOs (Fig. 1D and E). A subpopulation of GFP^+^ cells in the DFOs was colocalized with PAX6 staining (31.0 ± 1.6 % of total GFP^+^ cells), further indicating the dorsal forebrain identity of the OLIG2^+^ cells in the DFOs (Fig. 2E and F). Taken together, OLIG2 is not only expressed in the VFOs but also cytoplasmically expressed in a small subset of NPCs in the DFOs.

**Figure 2.**
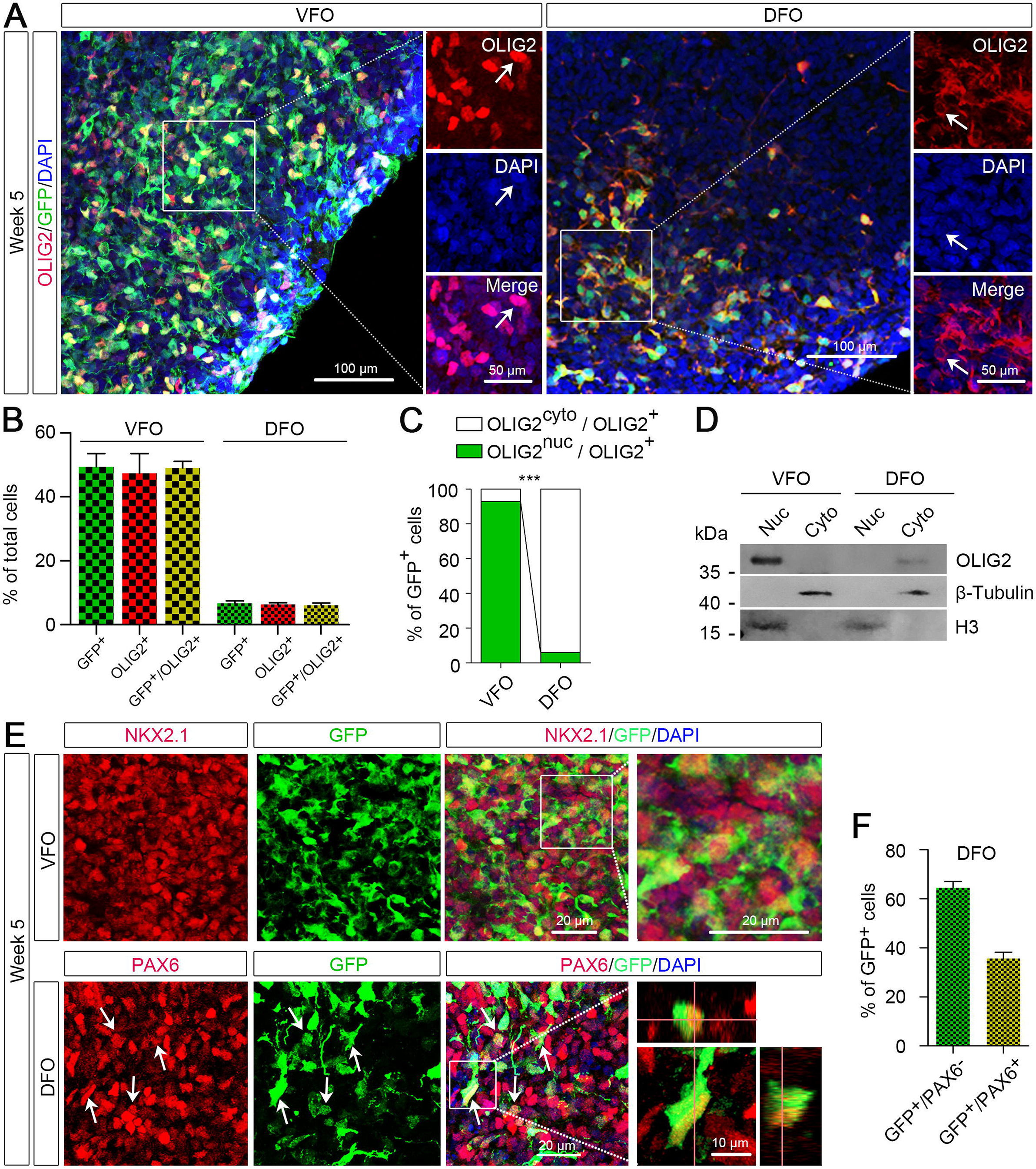
Cellular localization of OLIG2 in VFOs and DFOs. (A and B) Representatives and quantification of GFP^+^, OLIG2^+^, and GFP^+^/OLIG2^+^ cells in the VFOs and DFOs at week 5 (n = 4 organoids from two hPSC lines). Scale bars, 100 μm and 50 μm in the original and enlarged images, respectively. (C) Quantification of the percentage of GFP^+^ cells with nuclear or cytoplasmic OLIG2 expression among total GFP^+^ cells (n = 3 organoids from two hPSC lines). Student’s *t*-test., ***p < 0.001. (D) Western blot showing the cellular localization of OLIG2 extracted from week 5 organoids. The experiments were repeated for three times. For each experiment, at least 10 VFOs or DFOs derived from OLIG2-GFP hESCs and hiPSCs were pooled for protein extraction and fractionation. (E) Representative images of NKX2.1^+^/GFP^+^ cells in VFOs (top panels) and PAX6^+^/GFP^+^ cells in DFO (bottom panels). Scale bars, 50 μm and 10 μm in the original and enlarged images, respectively. (F) Quantification of percentage of GFP^+^/PAX6^−^ cells and GFP^+^/PAX6^+^ cells in week 5 DFOs (n = 5 organoids from two hPSC lines).

### OLIG2^+^ NPCs with dorsal forebrain regional identity can give rise to glutamatergic neurons

To further examine whether NPCs in the DFOs could differentiate into functional neurons and glial cells, we performed two-photon Ca^2+^ imaging on the organoids at week 6. We recorded patterns of Ca^2+^ transients from soma of total 78 spontaneously active cells collected from randomly selected 5 fields in 2 organoids. The Ca^2+^ transients exhibited by astrocytes are slower transients whereas the ones displayed by neurons are faster transients (Ohara et al., 2009; Tashiro et al., 2002). As shown in Fig. 3A and B, based on the duration and the number of oscillations, the patterns of Ca^2+^ transients could be grouped into two types: type 1 “glia-like” cells that had a lower number of oscillations with longer durations (> 10 s) and type 2 “neuron-like” cells that had a higher number of oscillations with shorter durations (< 2 s). Type 2 “neuron-like” cells also exhibited a significantly higher peak value of Ca^2+^ transients than type 1 “glia-like” cells (Fig. 3B).

**Figure 3.**
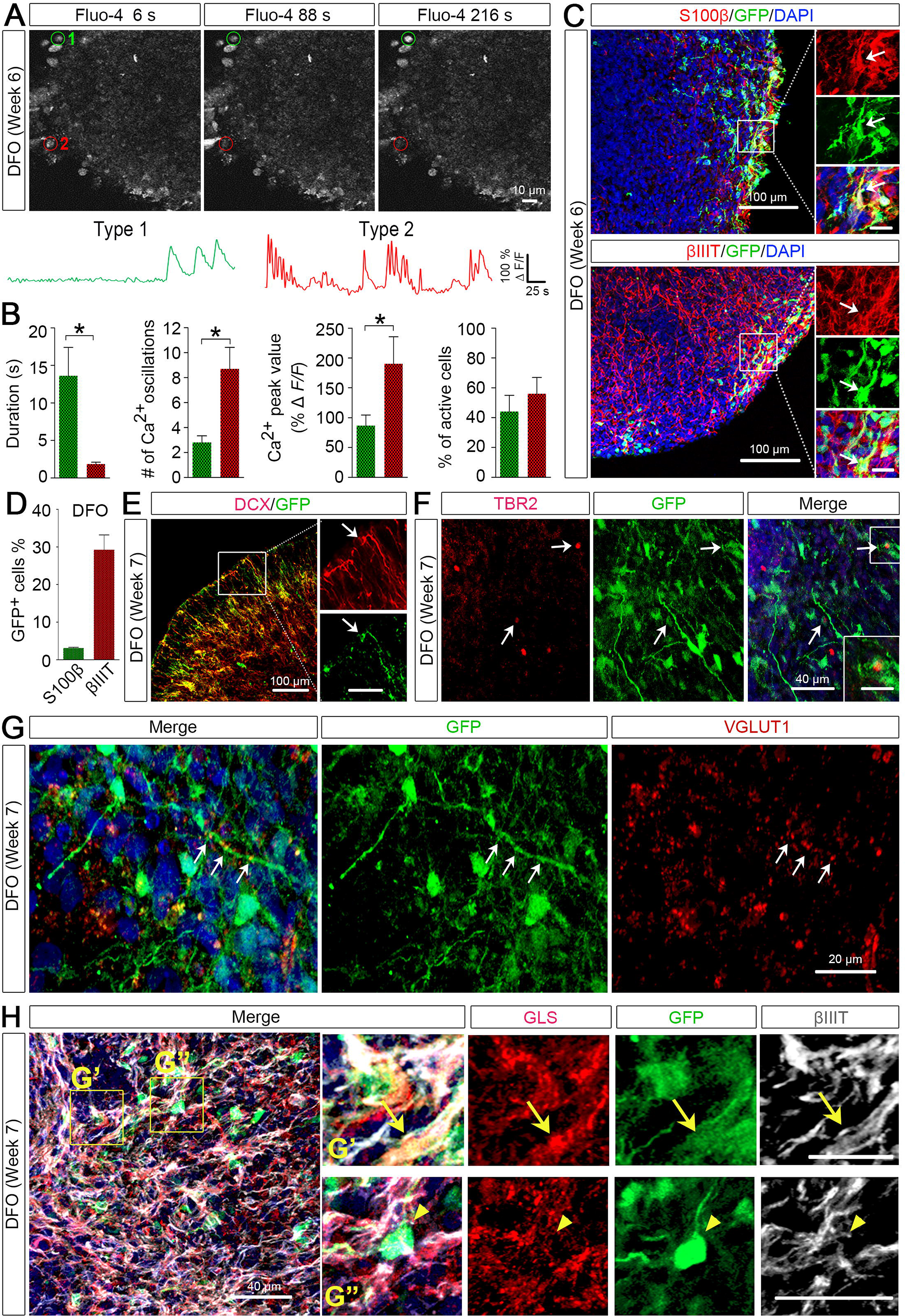
Neuronal differentiation of OLIG2^+^ cells in DFOs. (A) Time-lapse images and representative traces of Ca^2+^ transients in DFOs. ROIs of active Fluo-4^+^ cells were marked in red or green. (B) Quantification of Ca^2+^ transients. Scale bars represent 50 s and 100% ΔF/F. Quantification for the number, peak value, and duration of the Ca^2+^ transients (n = 5 fields from 2 organoids) are shown. (C and D) Representatives and quantification of GFP^+^/S100β^+^ and GFP^+^/βIIIT^+^ cells among total GFP^+^ cells (n = 3 organoids from two hPSC lines). Scale bars, 100 μm and 20 μm in the original and enlarged images, respectively. (E and F) Representative images showing co-localization of GFP with DCX or TBR2 in week 7 DFOs. Scale bars, 100 μm and 50 μm in the original and enlarged images for DCX. Scale bars, 40 μm and 20 μm in the original and enlarged images for TBR2. (G) Representative images showing co-localization of VGLUT1 and GFP. Scale bars, 20 μm. Arrows indicate VGLUT1 puncta along a GFP^+^ process. (H) Representative images showing co-localization of GLS, GFP, and βIIIT in week 7 DFOs. Arrows indicate a GFP^+^/βIIIT^+^/GLS^+^ neuron in the top panels of enlarged images, and arrowheads indicate a GFP^+^ cell that was GLS-negative in the bottom panels of enlarged images. Scale bars, 80 μm and 40 μm for top panels and bottom panels, respectively.

The nuclear localization of OLIG2 suggests differentiation to oligodendroglial lineage cells (Ligon et al., 2006), whereas cytoplasmic localization suggests differentiation to neuronal or astroglial cells (Setoguchi and Kondo, 2004; Takebayashi et al., 2000; Zhao et al., 2009). To delineate the identity of GFP^+^ cells, we immuno-stained the organoids with astroglial marker S100β or neuronal marker βIIIT. As shown in Fig. 3C, a large number of βIIIT^+^ neurons and a small population of S100β^+^ astrocytes were identified in 6-week-old DFOs. The GFP^+^ NPCs in DFOs mainly differentiated into neurons with a small percent giving rise to astroglia (Fig. 3D). In addition, we double-stained the DFOs with GFP and DCX, a marker for migrating immature neurons (Gleeson et al., 1999), or TBR2, a marker for intermediate neuronal progenitors (Sessa et al., 2010). There were GFP^+^/DCX^+^ processes and some GFP^+^ cells were positive for TBR2 (Fig. 3E), further suggesting that these GFP^+^/OLIG2^+^ cells were able to differentiate to neuronal cells.

We also found the formation of glutamatergic synapses in week 7 DFOs, as indicated by the staining of VGLUT1^+^ puncta (Fig. 3G). Notably, some of the VGLUT1^+^ puncta were found to distribute along the GFP^+^ processes. This prompted us to further characterize the OLIG2^+^ cells in the DFOs. We triple-stained the week 7 DFOs with GFP, βIIIT, and glutaminase (GLS) that is the enzyme essential for glutamate production in glutamatergic neurons and astrocytes in the brain (Aoki et al., 1991; Cardona et al., 2015). We found that a small population of GFP^+^ neurons, marked by βIIIT, expressed GLS (Fig. 3F), suggesting that the OLIG2^+^/GFP^+^ cells in the DFOs develop into glutamatergic neurons.

### BrainPhys medium promotes neuronal maturation in both VFOs and DFOs

Mounting evidence suggests that neuronal maturation and activity influence oligodendrogenesis and myelination (Gibson et al., 2014; Mitew et al., 2018). A recent study developed a new BrainPhys medium to better support the neurophysiological activity of cultured human neurons (Bardy et al., 2015). We further cultured DFOs and VFOs in the BrainPhys medium from week 7 to 9 (Fig. 4A). Then we examined the expression of c-Fos, an activity-dependent immediate early gene that is expressed in neurons following depolarization and often used as a marker for mapping neuronal activity (Loebrich and Nedivi, 2009). As shown in Fig. 4A and B, at week 9, there were significantly more c-Fos^+^ cells in the organoids cultured in BrainPhys medium than in ND medium. Furthermore, we observed that the synaptic markers, PSD-95 and SYNAPSIN 1 were also expressed in the DFOs cultured in BrainPhys medium (Fig. 4C). Moreover, neurons expressing CUX1, a marker for superficial-layer cortical neurons (Nieto et al., 2004) and TBR1^+^, a marker for pre-plate/deep-layer neurons (Hevner et al., 2001) were also seen in the DFOs (Fig. 4D). A small population of EMX1^+^ dorsal NPCs was also found in DFOs (Fig. 4E and F). At this time point, GFP signals became undetectable in the DFOs. In the VFOs, GFP fluorescence became much dimmer, compared to week 7 VFOs (Fig. 1G). This may result from the differentiation of a subset of OLIG2-expressing ventral forebrain NPCs to neurons, particularly interneurons (Xu et al., 2018), and subsequent termination of OLIG2/GFP expression in those cells, as reported in previous studies in mice (Miyoshi et al., 2007; Ono et al., 2008).

**Figure 4.**
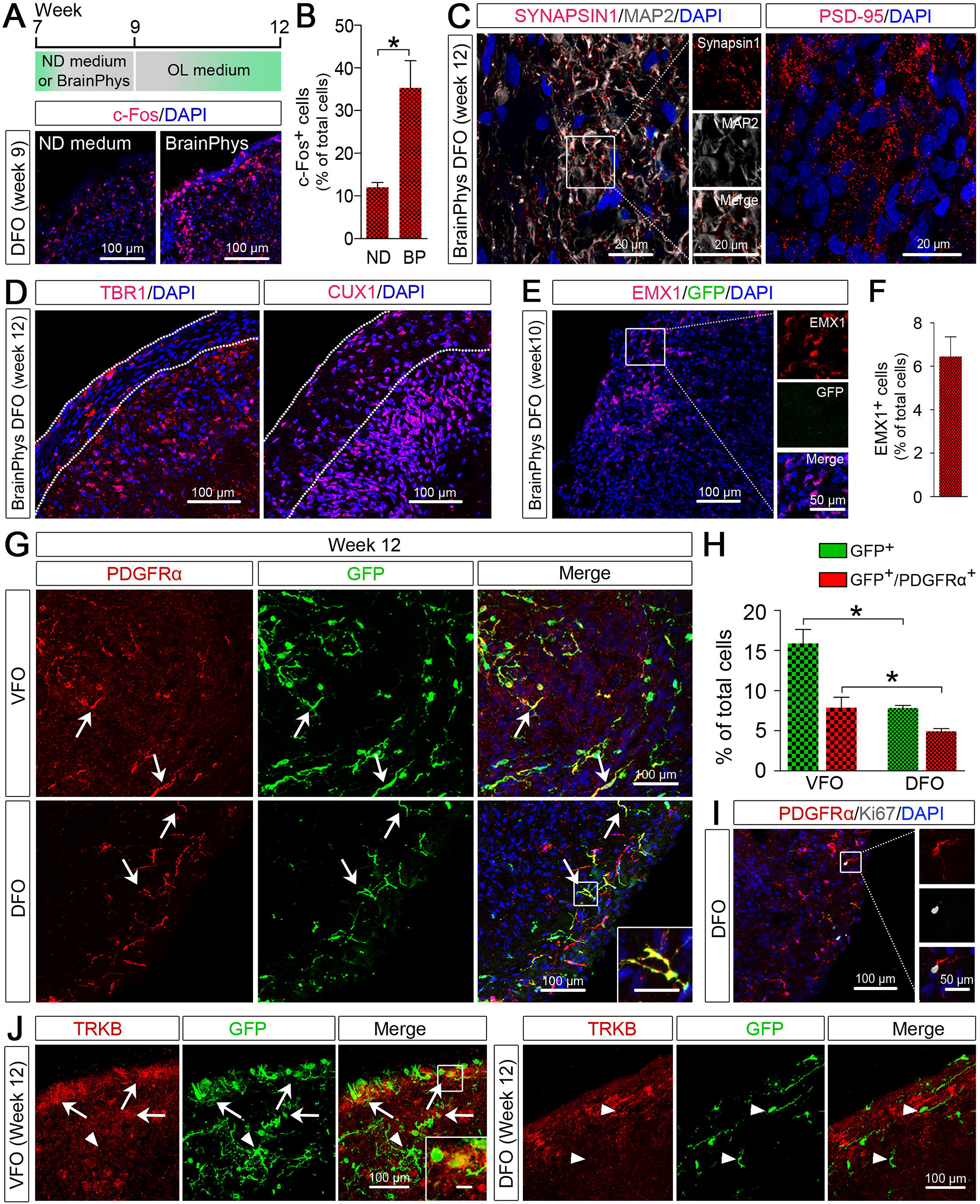
Oligodendrogenesis in VFOs and DFOs after neuronal maturation promoted by BrainPhys medium. (A and B) A schematic diagram, representatives, and quantification of c-Fos^+^ cells in week 9 DFOs cultured under different conditions (n = 4 organoids from two hPSC lines). Scale bar, 100 μm. Student’s *t*-test. *p < 0.05. (C) Representative images showing SYNAPSIN 1 and PSD-95 puncta in week 12 DFOs cultured in the BrainPhys medium. Scale bar, 20 μm. (D) Representative images showing TBR1^+^ and CUX1^+^ cells in week 12 DFOs cultured in BrainPhys medium. Scale bar, 100 μm. (E and F) Representatives and quantification of EMX1^+^ and GFP^+^ cells in week 10 DFO culture in BrainPhys medium (n = 4 organoids from two hPSC lines). Scale bars, 100 μm and 50 μm in the original and enlarged images, respectively. (G and H) Representatives and quantification of GFP^+^ and PDGFRα^+^ cells in week 12 VFOs and DFOs (n = 4 organoids from two hPSC lines). Arrows indicate the cells that are positive for PDGFRα and GFP. Scale, 100 μm. Student’s *t*-test. *p < 0.05. (I) Representatives of PDGFRα^+^/Ki67^+^ OPCs. Scale bars, 100 μm and 50 μm in the original and enlarged images, respectively. (J) Representatives of TRKB- and GFP-expressing cells in VFOs and DFOs. Arrows indicate TRKB^+^/GFP^+^ cells. Arrowheads indicate TRKB^−^/GFP^+^ cells. Scale bars, 100 μm and 20 μm in the original and enlarged images, respectively.

### Human PSC-derived organoids reveal oligodendrogenesis with ventral and dorsal origins

To promote oligodendrogenesis in the DFOs and VFOs, we further cultured the organoids for an additional 3 weeks (Fig. 4A; week 9 to 12) in OL differentiation medium containing thyroid hormone T3, which is commonly used to promote OL differentiation from hPSCs under 2D culture conditions (Goldman and Kuypers, 2015). As shown in Fig. 1G, at week 12, there were a large number of bright GFP^+^ cells evenly distributed in the VFOs. In the DFOs, GFP signal reappeared after the treatment of T3. To characterize the identity of these GFP^+^ cells, we double-stained GFP with OPC marker PDGFRα. In both VFOs and DFOs, nearly half of the GFP^+^ expressed PDGFRα, indicating the oligodendrogenesis in both organoids (Fig. 4G and H). A subset of PDGFRα^+^ OPCs in DFOs also exhibited immunoreactivity for Ki67, suggesting that these OPCs were capable of proliferation (Fig. 4I). In addition, we also stained the organoids with S100β or βIIIT, but found no GFP^+^ cells expressing S100β or βIIIT at week 12. Thus, the GFP^+^/PDGFRα-negative cells in the organoids might have committed to oligodendroglial fate but had not yet expressed any OPC marker. To further validate the ventral vs. dorsal origin of the OPCs in VFOs and DFOs, we examined the expression of tyrosine kinase B (TRKB). Previous studies (Du et al., 2003) have shown that TRKB is selectively expressed in oligodendroglia from the basal forebrain that has a ventral origin (Kessaris et al., 2006; Klambt, 2009), but not in oligodendroglia from cerebral cortex that has a dorsal origin (Kessaris et al., 2006; Winkler et al., 2018). TRKB was robustly expressed by neurons in both VFOs and DFOs. Only a small population of TRKB^+^/GFP^+^ cells were seen in VFOs but no TRKB^+^/GFP^+^ cells in DFOs (Fig. 4J). These results demonstrate that further culturing VFOs and DFOs under conditions favoring oligodendroglial differentiation can further reveal oligodendrogenesis with ventral and dorsal origins, respectively.

### OL maturation in DFOs, VFOs, and fused forebrain organoids (FFOs)

To examine whether OPCs in these organoids were able to mature into myelinating OLs, we cultured the VFOs and DFOs in OL medium for up to 6 weeks. Moreover, to test whether the interregional interactions between the differentially patterned human forebrain organoids are important for OL maturation, we generated FFOs and further cultured them in OL medium (Fig. 5A). VFOs at week 5 were used for fusion with DFOs because previous studies showed that immature migrating interneurons generated by ventral forebrain NPCs promote OL formation in the cortex in a paracrine fashion (Voronova et al., 2017). After fusion, we observed a massive migration of GFP^+^ cells from the VFOs into the DFOs (Fig. 5B), similar to the observations reported in previous studies (Bagley et al., 2017; Birey et al., 2017; Xiang et al., 2017). At 3 weeks after fusion (week 12), GFP^+^ cells largely populated the FFOs and exhibited bipolar morphology, characteristics of OPCs (Fig. 5B). At week 15, we assessed OL maturation in DFOs, VFOs, and FFOs by staining MBP, a marker for mature OLs. As shown in Fig. 5C and D, MBP^+^/GFP^+^ cells were detected in VFOs, but not in DFOs. Interestingly, fusing the organoids significantly promoted the maturation of OLs, as indicated by a higher percent of MBP^+^/GFP^+^ cells in FFOs. The mature OLs in FFOs exhibited complex processes at week 12 and occasionally, we observed tubular-shaped MBP staining in FFOs at week 15 (Fig. 5E), suggesting that the OLs started to myelinate axons. We further examined myelin ultrastructure in 8 sections from 4 FFOs. In total, we identified 52 myelinated axons. As shown in Fig. 5F, axons with loose escheatment by a few myelin laminae and some axons with compact myelin were observed in week 15 FFOs, which is similar to the “unorganized” myelin structure observed in oligocortical spheroids in a recent study (Madhavan et al., 2018) and also partly resembles the earliest stage of in vivo fetal myelinogenesis in humans (Weidenheim et al., 1992).

**Figure 5.**
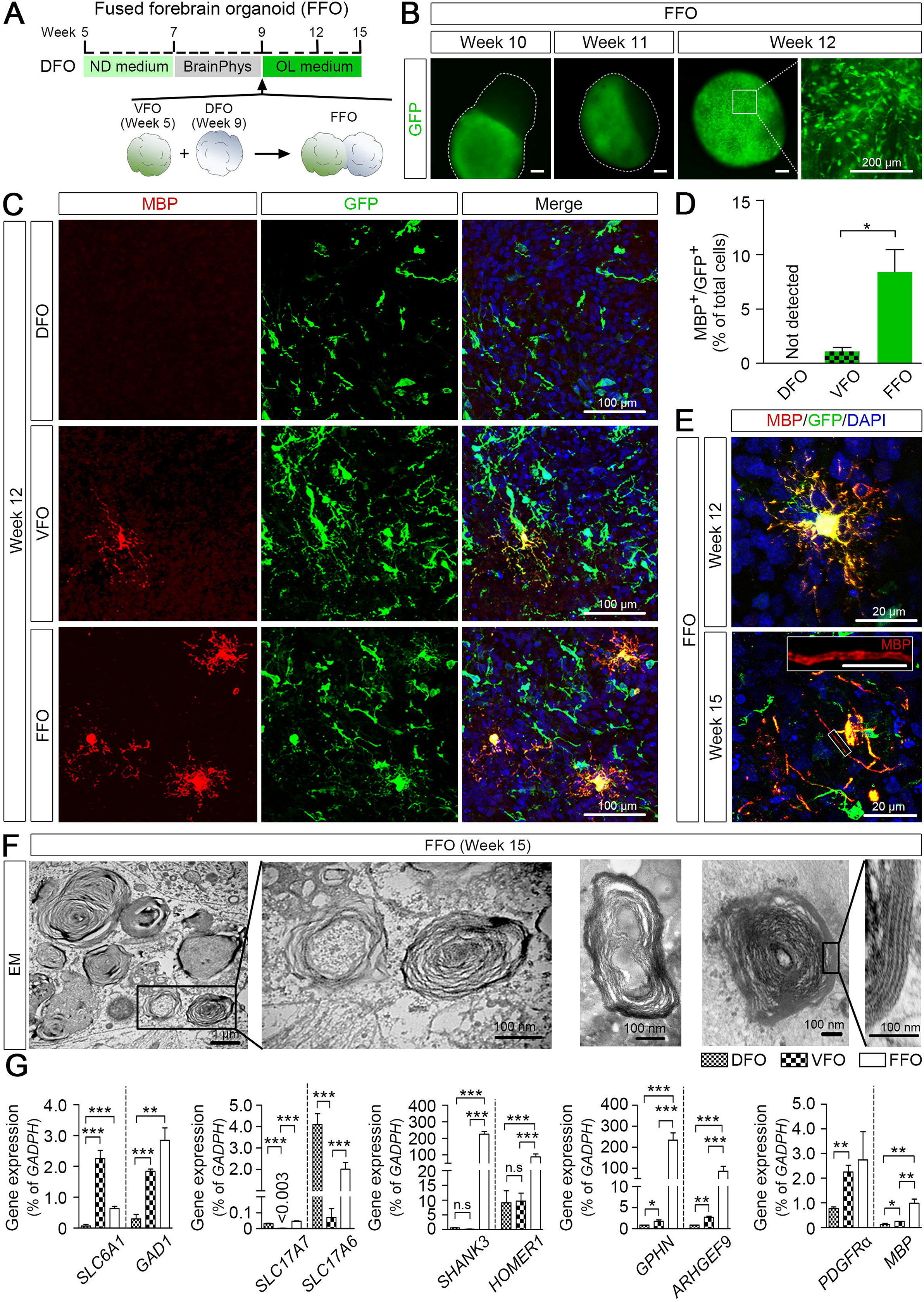
Oligodendroglial maturation in DFOs, VFOs, and FFOs. (A) A schematic procedure for fusing DFOs with VFOs and further culturing the FFOs in OL medium. (B) Representatives of GFP expression during the fusion process. FFOs are outlined with dotted lines at week 10 and 11. Scale bars, 200 μm. (C and D) Representatives and quantification of MBP^+^/GFP^+^ cells in VFOs, DFOs, and FFOs at week 12 (n = 4 organoids for each group). Scale bar, 100 μm. Student’s *t*-test. *p < 0.05. (E) Representatives of MBP^+^ cells in week 12 and week 15 FFOs. Scale bars, 20 μm and 10 μm in the original and enlarged images, respectively. (F) Representative EM images of the myelin ultrastructure in week 15 FFOs. Scale bars, 1 μm and 100 nm. (G) qRT-PCR results showing the expression of *SLC6A1*, *GAD1*, *SLC17A7*, *SLC17A6*, *SHANK3*, *HOMER1*, *GPHN*, *ARHGEF9*, *PDGFRα* and *MBP* in VFOs, DFOs, and FFOs (n = 3, independent experiments). One-way ANOVA with Tukey post-hoc test., ***p < 0.05, **p < 0.01, and ***p < 0.001.

We next examined the gene expression of markers for excitatory and inhibitory neurons as well as markers for inhibitory and excitatory synapses in VFOs, DFOs, and FFOs. As shown in Fig. 5G, the genes for markers of inhibitory neurons, including *SLC6A1* and *GAD1* that respectively encode GABA transporter type 1 and glutamate decarboxylase 1, were expressed at much higher levels in VFOs than in DFOs. In contrast, genes for markers of excitatory neurons, including *SLC17A7* and *SLC17A6* that respectively encode vesicular glutamate transporters VGLUT1 and VGLUT2, were expressed at higher levels in DFOs than in VFOs. Importantly, FFOs exhibited well-balanced expression of all these gene transcripts, as compared to DFOs and VFOs. FFOs exhibited not only significantly enhanced expression of markers for inhibitory neurons, compared to DFOs, but also enhanced expression of markers for excitatory neurons, compared to VFOs. Furthermore, we examined the expression of genes for components of post-synaptic machinery, such as *HOMER1* and *SHANK3* that respectively encode excitatory postsynaptic components HOMER and SHANK; and *ARHGEF9* and *GPHN* that respectively encode inhibitory postsynaptic components COLLYBISTIN and GEPHYRIN. Compared to DFOs and VFOs, FFOs exhibited significantly enhanced expression of the genes for components of both inhibitory and excitatory postsynaptic. The expression of genes for inhibitory postsynaptic components was higher in VFOs than in DFOs, and the expression of genes for excitatory postsynaptic components was similar in VFOs and DFOs. These findings demonstrate that the FFOs generated in our study possess integrated glutamatergic and GABAergic neurons, and have enhanced expression of synaptic markers, consistent with the results reported in recent studies (Birey et al., 2017; Xiang et al., 2017). Notably, the expression of *MBP* gene was also significantly higher in FFOs than in DFOs or VFOs. *PDGFRα* gene was expressed at a similar level in FFOs and VFOs, but at a lower level in DFOs. Our obervation suggests that enhanced neuronal network in our organoids might contribute to OL differentiation and maturation.

### Dorsally-derived oligodendroglia outcompete ventrally-derived oligodendroglia in the FFOs

To further examine which population of oligodendroglia constituted the oligodendroglial cells in FFOs, we generated a new version of fused forebrain organoids, by fusing the VFOs and DFOs that were respectively derived from isogenic OLIG2-GFP hiPSC reporter line and ND2.0 hiPSCs that do not have any reporter fluorescence (Fig. 6A). The isogenicity of OLIG2-GFP hiPSCs and ND2.0 hiPSCs were indicated by the identical short tandem repeat (STR) genotyping profile (Supplementary Table 1). In addition, we did not observe any growth advantage on one line versus the other between the two iPSC lines. Then we examined the interactions between ventrally- and dorsally-derived oligodendroglia in those FFOs. Since OLIG2 is expressed in all oligodendroglial lineage cells (Ligon et al., 2006), in this setting, dorsally- and ventrally-derived oligodendroglia could be readily distinguished because the former would be GFP-negative/OLIG2^+^, whereas the latter would be GFP^+^/OLIG2^+^. As shown in Fig. 6B and C, at one week after fusion (week 10), the ventrally-derived GFP^+^/OLIG2^+^ cells outnumbered the dorsally-derived GFP-negative/OLIG2^+^ cells. At 3 and 6 weeks after fusion (week 12 and 15), the dorsally-derived GFP-negative/OLIG2^+^ cells became a dominant population, whereas the ventrally-derived GFP^+^/OLIG2^+^ cells dramatically decreased in the FFOs from week 10 to 15 (Fig. 6B and C). Furthermore, at week 12, the majority of MBP^+^ cells were MBP^+^/GFP-negative (Fig. 6D and E), suggesting that those mature OLs had a dorsal origin. Taken together, these results demonstrate that the majority of oligodendroglia in the FFO at an early stage (week 10) are ventrally-derived oligodendroglia, which are outcompeted by dorsally-derived oligodendroglia in the FFOs at later stages (week 12 and 15).

**Figure 6.**
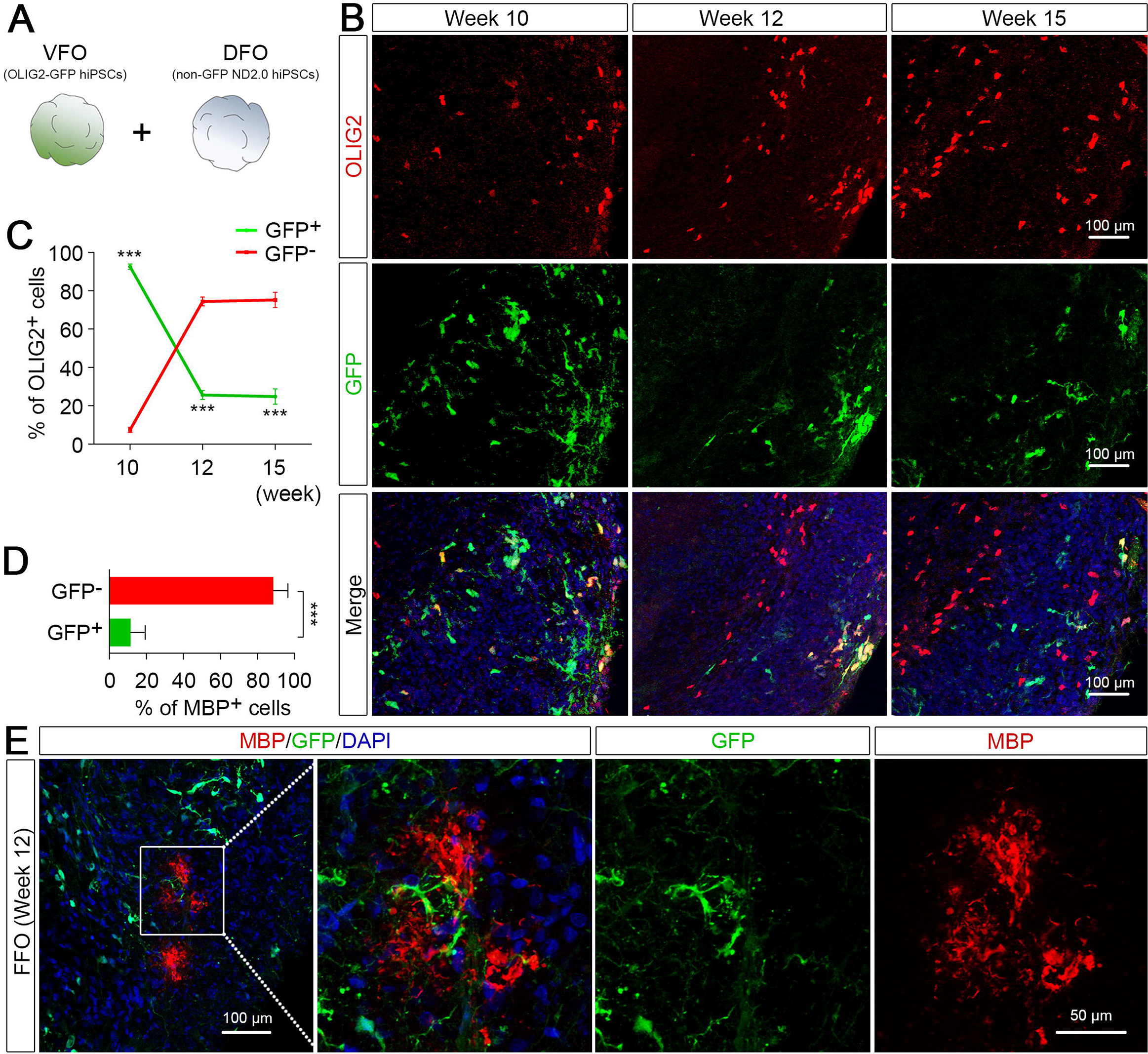
Interactions between dorsally- and ventrally-derived oligodendroglia in the FFOs. (A) A schematic procedure for fusing the VFOs and DFOs that were respectively derived from isogenic OLIG2-GFP hiPSC reporter line and ND2.0 hiPSC line that do not have any reporter fluorescence. (B and C) Representatives and quantification (n = 4 organoids) showing that ventrally-derived OLIG2^+^ cells (GFP^+^) are outnumbered by dorsally-derived OLIG2^+^ cells (GFP-negative) at one week after fusion (week 10) to six weeks after fusion (week 15). Scale bar, 100 μm. Student’s *t*-test., ***p < 0.001. (D and E) Representatives and quantification showing the MBP^+^ cells with a ventral (GFP^+^) or dorsal (GFP-negative) origin (n = 4 organoids). Scale bars, 50 μm. Student’s *t*-test., ***p < 0.001.

## DISCUSSION

OLIG2 is not only closely associated with the development of OLs in the vertebrate CNS, but also is expressed in NPCs during development (Jiang et al., 2013; Ligon et al., 2006; Meijer et al., 2012; Ono et al., 2008). At early embryonic stages in mice, OLIG2 is mostly expressed in the brain by the NPCs that distribute in the ventral telencephalon (Ligon et al., 2006; Meijer et al., 2012). A distinct but small number of OLIG2^+^ NPCs that distribute in the dorsal forebrain has been also identified in both mouse and human embryonic brain (Jakovcevski and Zecevic, 2005; Ono et al., 2008). The small population of human OLIG2^+^/PAX6^+^ NPCs seen in the week 5 DFOs may mimic the OLIG2^+^ NPCs in the human embryonic diencephalon (Jakovcevski and Zecevic, 2005). Furthermore, we demonstrate that these early human OLIG2^+^ NPCs are also able to give rise to glutamatergic neurons and astrocytes in the organoids, similar to that seen in the mouse brain (Ono et al., 2008).

Interestingly, we revealed oligodendrogenesis with a dorsal origin in our organoid models. This oligodendrogenesis in DFOs likely recapitulates the 3^rd^ wave of oligodendrogenesis because: First, these oligodendroglial cells are dorsally-derived. The DFOs are generated by using primitive NPCs (pNPCs) as the starting population, since these neural fate-restricted pNPCs retain high responsiveness to instructive neural patterning cues (Li et al., 2011) and can be efficiently patterned to desired brain regions (Monzel et al., 2017). We further enhanced the dorsalization using CycA (Bagley et al., 2017; Vazin et al., 2014). In addition, we did not observe any ventral forebrain NPCs in DFOs. Second, these oligodendroglial cells are derived from progenitors that are distinct from the early OLIG2^+^/PAX6^+^ progenitor cells. At week 9, all OLIG2^+^/PAX6^+^ NPCs differentiate to neurons or astrocytes, and OLIG2 expression terminates as indicated by GFP fluorescence and qPCR results. Moreover, we find EMX1^+^ dorsal forebrain NPCs in the 9-week-old organoids. It is very likely that those EMX1^+^ cells further differentiate to oligodendroglial cells, similar to the findings in the mouse brain (Kessaris et al., 2006; Winkler et al., 2018). Third, the newly emerged OLIG2^+^/PDGFRα^+^ cells in week 12 DFOs are able to proliferate and populate the organoids, indicating their OPC nature. Lastly, these dorsally-derived oligodendroglia can outcompete ventrally-derived oligodendroglia in the FFOs, which may mimic the process of oligodendrogenesis in the cerebral cortex, where ventrally derived oligodendroglia are outcompeted by oligodendroglia with a dorsal origin at late developmental stages (Kessaris et al., 2006). Thus, these results suggest that a dorsal origin of oligodendrogenesis may occur during human brain development.

Technically, we also develop organoid models with significantly accelerated OL maturation. Deriving mature OLs from human PSCs is a tedious and lengthy process, and optimizing methods to obtain human mature OLs has been a major research effort (Douvaras and Fossati, 2015; Goldman and Kuypers, 2015; Liu et al., 2011; Stacpoole et al., 2013). In our system, MBP^+^ mature OLs appeared at 9 weeks after organoid formation. This acceleration of the OL maturation program in our organoids is likely achieved by promoting neuronal maturation with the newly-designed BrainPhys medium and rebuilding the neuronal network in FFOs. Balanced excitatory and inhibitory neuronal activities (Nagy et al., 2017), neurotransmitters and neurotrophins released by excitatory neurons or astrocytes (Gautier et al., 2015; Lundgaard et al., 2013; Xiao et al., 2009), as well as cytokines released by young migrating inhibitory neurons (Voronova et al., 2017) may partly contribute to the enhanced oligodendroglial lineage progression in FFOs. As compared to the fusion methods, generating a single, regionalized organoids containing both ventral and dorsal elements (Cederquist et al., 2019) may more precisely recapitulate the serial waves of oligodendroglial production and replacement seen in the developing forebrain, because the different germinal zones within single organoids might be well preserved and not be disrupted by the fusion process. Notably, similar to the recent studies (Madhavan et al., 2018; Marton et al., 2019), the maturation of human OPCs is not efficient in organoids, and those myelinated axons often show wrapping with multiple layers of uncompacted myelin. This may be attributed in part to a lack of oxygen penetration in the organoids cultured for long-term, which can result in necrosis in the core of organoids (Brawner et al., 2017; Giandomenico and Lancaster, 2017). As opposed to rodent OPCs, human OPC differentiation and myelination are highly sensitive to hypoxic conditions (Gautier et al., 2015). Future studies aiming at integrating a vascular structure that brings the adequate delivery of oxygen and nutrients and promoting neuronal maturation and electrical activity (Brawner et al., 2017; Mansour et al., 2018) may help facilitate organoid models with robust OL maturation.

Previous *in vivo* studies have helped us understand the developmental differences between ventrally- and dorsally-derived OPCs (Kessaris et al., 2006; Nery et al., 2001; Orentas et al., 1999; Winkler et al., 2018). However, it is largely unclear what the phenotypic and functional differences are between the different populations of OPCs. In this study, we demonstrate the generation of dorsally- and ventrally-derived human oligodendroglial cells in organoids. We propose that the competitive advantage exhibited by dorsally-derived over ventrally-derived OPCs may partly result from their unique expression pattern of molecules involved in controlling OPC proliferation and lineage progression, for example GPR17, a G-protein-coupled membrane receptor suggested as an intrinsic timer of OL differentiation during development (Chen et al., 2009; Fumagalli et al., 2011), and transcription factor EB (TFEB), which recently has been identified to govern the regional and temporal specificity of OL differentiation and myelination (Sun et al., 2018). Future gene expression profiling at a single-cell level in the separate dorsal and ventral forebrain organoids as well as FFOs will significantly advance our understanding on the heterogeneity of human OPCs. Combing with human iPSC technologies, examining oligodendroglia in region-specific organoids may be more informative to understand disease mechanisms of neurodevelopmental disorders associated with myelin defects. These organoid models could also have important implications in the setting of CNS injury, as distinct populations of human OPCs might respond to pro-myelinating drugs differently, preferentially contributing to remyelination, thus enabling a better model for therapeutic manipulation.

## EXPERIMENTAL PROCEDURES

See further details in the Supplemental Experimental Procedures.

### Generation of human forebrain organoids with hPSC lines

All the hPSC studies were approved by the committees on stem cell research at Rutgers University. Forebrain organoids differentiation was carried out as previously reported (Bagley et al., 2017; Birey et al., 2017; Xiang et al., 2017) with modifications.

### qRT-PCR

Gene expressions were measured by performing qRT-PCR with TaqMan primers listed in table S2.

### Western blotting

Protein expression and localization in cells were evaluated by immunoblotting with samples achieved by REAP method (Suzuki et al., 2010).

### Immunostaining

Fixed organoids were processed for Immunostaining with primary antibodies listed in Table S3.

### Calcium imaging

DFOs loaded with fluo-4 AM (5 μM, Molecular Probes) was performed with a two-photon microscope (Moving Objective Microscope; Sutter Instruments).

### Electron microscopy

Selected vibratome sections from organoid samples were used for electron microscopy.

### DNA fingerprinting short tandem repeat (STR) analysis

GENEprint PowerPlex 16 kit (Promega performed by Cell Line Genetics. LLC) was used for STR analysis.

### Statistical analysis

All experiments were repeated at least three times (n ≥ 3). Unless otherwise noted, organoids derived from OLIG2-GFP hESCs and hiPSCs were analyzed for each experiment. Data are presented as mean ± S.E.M. and were analyzed using one-way ANOVA in combination with Tukey post-hoc test or Student’s *t*-test. For statistical significance, p values < 0.05 were considered significant.

## Acknowledgments

This work was in part supported by grants from the NIH (R21HD091512 and R01NS102382 to P.J.) and a pilot grant from Rutgers Brain Health Institute to P.J.

## Author Contributions

H.K. and P.J. designed experiments and interpreted data; H.K. carried out most of experiments with technical assistance from R.X.; P.R., and A.D. performed calcium imaging experiments; Y.L. and C.D. provided study materials and reagents and provided critical suggestions to the overall research direction. P.J. directed the project and wrote the manuscript together with H.K. and input from all co-authors.

