## supplemental info for "Pluripotent Stem Cell-derived Cerebral Organoids Reveal Human Oligodendrogenesis with Dorsal and Ventral Origins"

### Supplemental Experimental Procedures

#### Culture and derivation of hPSC cell lines

The OLIG2-GFP knockin hPSCs (hESCs and hiPSCs) were established using a gene-targeting protocol and fully characterized, as reported in our previous studies (Liu et al., 2011; Xue et al., 2016). The OLIG2-GFP hiPSC reporter line was established from the ND2.0 hiPSC line that was obtained from Center for Regenerative Medicine, National Institutes of Health. The hPSCs maintained under a feeder-free condition on a hESC-qualified Matrigel (Corning)-coated dish with mTeSR1 media (STEMCELL Technologies) were used for this study. The hPSCs were passaged approximately once per week by ReLeSR media (STEMCELL Technologies).

#### Generation of human forebrain organoids

To avoid non-CNS differentiated tissue and reduce variability in 3D cerebral organoids generation, we used purified pNPCs as the starting population for generating organoids. As shown in Fig. 1A, we induced pNPCs from hPSCs using a small molecule-based protocol (Chen et al., 2016; Li et al., 2011). Briefly, neural differentiation was induced by dual inhibition of SMAD signaling (Chambers et al., 2009) with inhibitors SB431542 (5 μM, Stemgent) and noggin (50 ng/ml, Peprotech) for a week. The embryoid bodies (EB) were then plated on dishes coated with growth factor-reduced Matrigel (BD Biosciences) in the medium consisting of DMEM/F12, 1x N2, and laminin (1 μg/ml; Sigma-Aldrich) for a week. Next, neural rosettes were manually isolated from the expanded area. The isolated neural rosettes were further cultured in pNPC media, composed of a 1:1 mixture of Neurobasal (Thermo Fisher Scientific) and DMEM/F12, supplemented with 1 x N2, 1 x B27-RA (Thermo Fisher Scientific), FGF2 (20 ng/ml, Peprotech), human leukemia inhibitory factor (hLIF, 10 ng/ml, Millipore), CHIR99021 (3 μM, Stemgent), SB431542 (2 μM), and ROCK inhibitor Y-27632 (10 μM, Tocris). To generate organoids, dissociated pNPCs by TrypLE Express (Thermo Fisher Scientific) were placed into low-attachment 96-well plates at a density of 9,000 cells to develop uniform organoids for two days. The pNPC aggregates were then grown and patterned in low-attachment 6-well plates with the treatment of either 5 μM Cyclopamine A (Cyc A; Calbiochem) for dorsalization or dual activation of SHH pathway with sonic hedgehog (SHH; 50 ng/ml, Peprotech) and purmorphamine (Pur; 1 μM, Cayman Chem) for ventralization. Starting from week 5, the DFOs were cultured on an orbital shaker with a speed of 80 rpm/min in neuronal differentiation (ND) medium containing a 1:1 mixture of Neurobasal and DMEM/F12, supplemented with 1 x N2, 1 x B27, BDNF (20 ng/ml, Peptrotech), GDNF (20 ng/ml, Peprotech), dibutyryl-cyclic AMP (1mM, Sigma), and ascorbic acid (200 nM, Sigma). The 5-week-old VFOs were cultured in OPC medium containing DMEM/F12, supplemented with 1 x N2, 1 x B27, FGF2 (10 ng/ml, Peptrotech), PDGF-AA (10 ng/ml, Peprotech). For further neuronal maturation, both DFOs and VFOs were cultured in BrainPhys medium (STEMCELL Technologies). Starting from week 9, organoids were maintained in ND medium supplemented with 3,3,5-Triiodo-L-thyronine sodium salt (T3; 10 ng/ml, Cayman Chem; OL medium) for oligodendroglial differentiation and maturation. FFOs were generated by using a spontaneous fusion method (Bagley et al., 2017; Birey et al., 2017; Xiang et al., 2017) with modifications. Briefly, single week 9 DFO were closely placed with a week 5 VFO by transferring both of them into the round-bottom ultra-low-attachment 96-well plate for 2 days without agitating. Then, the FFOs were transferred to an ultra-low-attachment 6-well plate and cultured for a day without agitating. The next day, the FFOs were maintained with OL medium on an orbital shaker with a speed of 80 rpm/min.

**RNA isolation and qRT-PCR**

Total RNA extracted from organoids with RNAeasy kit (Qiagen) was used to make complementary DNA with a Superscript III First-Strand kit (Invitrogen). The qRT-PCR was performed with TaqMan primers listed in supplementary table 2 on an Abi 7500 Real-Time PCR system. Experimental samples were analyzed by normalization with the expression level of housekeeping gene glyceraldehyde-3-phosphate dehydrogenase (GAPDH). Relative quantification was performed by applying the $2^{-\Delta\Delta Ct}$ method (Livak and Schmittgen, 2001).

**Western blotting**

OLIG2 protein expression and localization in cells were evaluated by immunoblotting using fractionated samples. The nuclear and cytoplasmic fraction was achieved by modification of a reported method (Suzuki et al., 2010). Briefly, organoids were washed with PBS and harvested. Resuspended organoids in 900 ul of ice-cold lysis buffer (0.1% NP40 in PBS) were lysed 10 times through a 25-gauge syringe. Following the second spinning down for 30 seconds, the supernatant was collected as a cytosolic fraction. 3 times washed pellet was lysed in sample buffer containing 1% of SDS. Fractionated proteins were separated on 12% SDS-PAGE gel and transferred onto nitrocellulose membrane. Blots were then blocked in 2% skim milk and incubated with primary antibodies at 4°C overnight. The information for primary antibodies and dilutions is listed in Supplementary table 3. Afterward, the blots were incubated with secondary antibodies conjugated with a fluorophore for an hour at room temperature. Western blot was visualized using Odyssey (LiCor).

**Immunostaining and cell counting**

Organoids fixed with 4% paraformaldehyde were processed and cryo-sectioned for immunofluorescence staining. The information for primary antibodies and dilutions is listed in Table S2. Slides were mounted with the anti-fade Fluoromount-G medium containing 1,40,6-diamidino-2-phenylindole dihydrochloride (DAPI) (Southern Biotechnology). Images were captured with LSM800 confocal microscope. The cells were counted with ImageJ software. At least six fields chosen randomly from three sections of each organoid were counted. For each organoid, at least 400 cells were counted.

**Dye loading and calcium imaging**

For calcium imaging in organoids, DFOs were placed on the growth factor-reduced Matrigel-coated coverslip in 6-well plate for 24 hr. The next day, organoids were loaded with fluo-4 AM (5 μM, Molecular Probes) and 0.04% Pluronic F-127 for 40 minutes and then transferred to the neuronal differentiation (ND) medium for at least 30 minutes before transferring to the submersion-type recording chamber (Warner) superfused at room temperature with artificial CSF (ACSF, mM: 126 NaCl, 3 KCl, 1.25 $NaH_2PO_4$, 1 $MgSO_4$, 2 $CaCl_2$, 26 $NaHCO_3$, and 10 dextrose) saturated with 95% $O_2$/5% $CO_2$. All imaging was performed with a two-photon microscope (Moving Objective Microscope; Sutter Instruments) coupled to a Ti:Sapphire laser (Chameleon Vision II, Coherent). Images were collected with a Nikon water immersion objective (25X, 1.05 NA). Excitation power measured at the back aperture of the objective was typically about 20-30 mW, and laser power was modulated using a Pockels cell. Fluo-4 was excited at 820nm and emission was detected with GaAsP detector

(Hamamatsu Photonics) fitted with a 535/50 bandpass filter and separated by a 565 nm dichroic mirror. ScanImage (v5.1, Vidrio Technologies) software (Pologruto et al., 2003) was used for imaging. Time-lapse imaging was performed every 1 s for 5 minutes and was collected at 512 by 512-pixel resolution. Image analysis was performed using ImageJ software. Regions of interest (ROIs) were placed around soma. Fluorescence was averaged over ROIs placed and expressed as relative fluorescence changes ($\Delta F/F$) after subtraction of background fluorescence from a neighboring region. For each ROI, basal fluorescence was determined during 20 s periods with no $Ca^{2+}$ fluctuation. $Ca^{2+}$ transients were detected when fluorescence intensity reached higher than 2 SD value of baseline fluorescence intensity.

**Electron microscopy**

Organoid samples were fixed with 2% glutaraldehyde, 2% paraformaldehyde, and 0.1M sodium cacodylate in PBS. The selected vibratome sections were post-fixed and processed for electron microscopy (EM) as described in our previous studies (Chen et al., 2016; Jiang et al., 2016). EM images were captured using a high-resolution charge-coupled device (CCD) camera (FEI).

**DNA fingerprinting short tandem repeat (STR) analysis**

STR analysis was performed using GENEprint PowerPlex 16 kit (Promega performed by Cell Line Genetics. LLC). Samples were run in duplicate and blinded to the interpreter to confirm the results. Please note that ND2.0 and OLIG2-GFP hiPSC lines have identical STR genotyping profile, indicating that they are isogenic lines derived from the same parental cell line.

**Supplementary Table 1. STR genotyping profile of ND2.0 hiPSCs and OLIG2-GFP hiPSC reporter line.** STRs of all loci for OLIG2-GFP hiPSCs match to ND2.0 hiPSCs.

| STR Locus | Chr. Location | ND2.0 hiPSCs | | OLIG2-GFP hiPSCs | |
|---|---|---|---|---|---|
| **Amelogenin** | **Xp22.1-22.3 and Y** | **X** | **Y** | **X** | **Y** |
| vWA | 12p12-pter | 17 | 18 | 17 | 18 |
| D8S1179 | 8q | 12 | | 12 | |
| TPOX | 2p23-2pter | 10 | 11 | 10 | 11 |
| FGA | 4q28 | 24 | 26 | 24 | 26 |
| D3S1358 | 3p | 15 | | 15 | |
| THO1 | 11p15.5 | 6 | 9.3 | 6 | 9.3 |
| D21S11 | 21q11-21q21 | 29 | 31.2 | 29 | 31.2 |
| D18S51 | 18q21.3 | 13 | 18 | 13 | 18 |
| Penta E | 15q | 7 | 12 | 7 | 12 |
| D5S818 | 5q23.3-32 | 11 | 12 | 11 | 12 |
| D13S317 | 13q22-q31 | 11 | 12 | 11 | 12 |
| D7S820 | 7q11.21-22 | 12 | | 12 | |
| D16S539 | 15q24-qter | 9 | 11 | 9 | 11 |
| CSF1PO | 5q33.3-34 | 12 | 13 | 12 | 13 |
| Penta D | 21q | 9 | 15 | 9 | 15 |

**Supplementary Table 2. A list of primers used.**

| Gene | Gene expression assay catalog number |
|---|---|
| *ARHGEF9* | HS01003480_m1 |
| *DLX1* | Hs00269993_m1 |
| *EMX1* | Hs00417957_m1 |
| *GAD1* | Hs01065893_m1 |
| *GAPDH* | Hs02758991_g1 |
| *GPHN* | HS00982840_m1 |
| *HIF1A* | Hs00153153_m1 |
| *HOMER1* | Hs01029333_m1 |
| *LEF1* | Hs01547250_m1 |
| *LHX6* | Hs01030941_g1 |
| *MBP* | Hs00921945_m1 |
| *NKX-2-2* | Hs05035641_s1 |
| *OLIG2* | Hs00300164_s1 |
| *PDGFRα* | Hs00998018_m1 |
| *S100β* | Hs00389217_m1 |
| *SHANK3* | Hs01393541_m1 |
| *SLC17A6 (VGLUT2)* | Hs00220439_m1 |
| *SLC17A7 (VGLUT1)* | Hs00220404_m1 |
| *SLC6A1 (GAT1)* | Hs01104475_m1 |
| *TBR2* | Hs00232429_m1 |

**Supplementary Table 3. A list of antibodies used.**

| Antibodies | Vendor/Catalog #. | Type | Dilution |
|---|---|---|---|
| βIIItubulin | Millipore / MAB1637 | Mouse IgG | 1:200 |
| β-tubulin | DSHB / E7 | Mouse IgG | WB (1:1000) |
| c-FOS | Santa Cruz / SC-52 | Rabbit IgG | 1:100 |
| CUX1 | Santa Cruz / SC13024 | Rabbit IgG | 1:500 |
| DCX | Cell Signaling / 4604s | Rabbit IgG | 1:500 |
| EMX1 | Sigma / HPA006421 | Rabbit IgG | 1:1000 |
| FOXG1 | Abcam / ab18259 | Rabbit IgG | 1:500 |
| GFAP | Millipore / AB5804 | Rabbit IgG | 1:1000 |
| GFP | Rockland / 600-141-215 | Goat IgG | 1:1000 |
| GFP | Thermo / MA5-15256 | Mouse IgG | 1:500 |
| GLS | Abcam / ab156876 | Rabbit IgG | 1:250 |
| Ki67 | Cell signaling / 9449 | Mouse IgG | 1:400 |
| Ki67 | Thermo Fisher Scientific / SP6 | Rabbit IgG | 1:200 |
| MAP2 | Millipore / AB3418 | Mouse IgG | 1:500 |
| LHX6 | Abcam / ab22885 | Rabbit IgG | 1:100 |
| MBP | Millipore / MAB386 | Rat IgG | 1:100 |
| NeuN | Millipore / MAB377 | Mouse IgG1 | 1:100 |
| Nestin | Santa Cruz / SC-21249 | Goat IgG | 1:100 |
| NKX2.1(TTF1) | Abcam / ab76013 | Rabbit IgG | 1:200 |
| OLIG2 | Phosphosolutions 1538 | Rabbit IgG | 1:1000; WB (1:2000) |
| PAX6 | GeneTex / GTX11324 | Rabbit IgG | 1:400 |
| PDGFRα | Santa Cruz / SC338 | Rabbit IgG | 1:50 |
| P-Histone H3 | Thermo Fisher Scientific / PA5–17869 | Rabbit IgG | WB(1:1000) |
| PSD95 | Invitrogen 51-6900 | Rabbit IgG | 1:100 |
| S100β | Sigma / S2532 | Mouse IgG | 1:1000 |
| SOX2 | Millipore / AB5603 | Rabbit IgG | 1:100 |
| Synapsin I | Millipore / AB1543P | Rabbit IgG | 1:400 |
| TBR1 | EMD Millipore / AB2261 | Chicken IgG | 1:100 |
| TBR2 | Abcam / AB23345 | Rabbit IgG | 1:100 |
| VGLUT1 | Millipore / AB5905 | Guinea pig IgG | 1:250 |

Antibody dilution for western blotting are specifically marked as WB, and others are for immunostaining.
